# The dependence of the amino acid backbone conformation on the translated synonymous codon is not statistically significant

**DOI:** 10.1101/2022.11.29.518303

**Authors:** Javier González-Delgado, Pablo Mier, Pau Bernadó, Pierre Neuvial, Juan Cortés

## Abstract

In their recent work, Rosenberg *et al*. [1] studied the dependence between the identity of synonymous codons and the distribution of the backbone dihedral angles of the translated amino acids. In the past, it has been shown that the use of synonymous codons is highly relevant in multiple biological processes including, among others, mRNA splicing, translational rates and protein folding [2, 3]. While the correlation between synonymous codons and secondary structure in translated proteins has been widely studied [4–6], Rosenberg *et al*. evaluated the effect of codon identity on a finer scale, analyzing whether the distribution of (*ϕ, ψ*) dihedral angles within secondary structure elements is significantly altered when synonymous codons are used. Their conclusion, showing significant differences, particularly for amino acid residues involved in *β*-strands, would represent a new paradigm for the role played by synonymous codons in defining protein structure. However, the statistical methodology used in that study was formally incorrect, casting doubt on the obtained results. Besides, it is based on density estimates that might be imprecise for small sample sizes, yielding misleading comparisons. These methodological errors are described in the following section. Then, using an appropriate methodology, we reanalyzed the data presented in [1]. Our results show that the influence of the codon on the distribution of the dihedral angles is statistically non-significant for all types of secondary structures, contradicting the conclusion by Rosenberg *et al*.. These results were corroborated by repeating the analysis on structures extracted from the AlphaFold Database [7, 8] for the same set of proteins, and shown to be robust with respect to the definition secondary structural classes and also when considering the nature of the neighbor residues. Overall, our observations demonstrate that the influence of the synonymous codons on the backbone dihedral angles can not be inferred with current data.

## Limitations of the original methodology

### Ill-defined p-values

The goal of Rosenberg *et al*. was to assess the effect of synonymous codons on the distribution of (*ϕ, ψ*) dihedral angles by comparing codon-specific Ramachandran plots. Keeping the notation of [1], if (*c, c′*) denotes a pair of synonymous codons and *𝒳* a type of secondary structure, they aimed at testing the null hypothesis *H*_0,(*c,c′)*|*𝒳*_ that both codon-specific distributions are the same. To do so, the authors introduced a metric to quantify differences between the distributions corresponding to different codons. Then, to assess the significance of such differences, Rosenberg *et al*. proposed to draw *B* = 25 pairs of bootstrapped samples, and to compare them with their synonymous codon counterparts using a permutation test procedure, with *K* = 200 permutations. For each bootstrap sample *b* ∈ {1, *…, B*}, if *n*_*b*_ denotes the number of permutations where the permuted metric is larger than the base metric (obtained from non-permuted data), they proposed the quantity

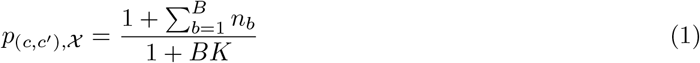

as a *p*-value for *H*_0,(*c,c′)*|*𝒳*_. We can reformulate (1) in order to gain insight into its statistical behavior. First, let us define

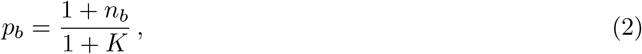

which is a well-defined *p*-value for the *b*-th permutation test. Letting

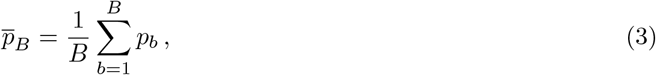

it can be shown (see SI) that

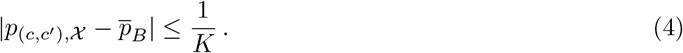

That is, for sufficiently large *K, p*_(*c,c′),𝒳*_ is approximately the empirical mean of the *B p*-values associated to individual permutation tests.

However, 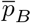 is not a valid *p*-value (see SI for a formal proof). Let us recall that a *p*-value *p* is statistically valid if and only if its distribution under the null hypothesis is Super-Uniform. A random variable is said to be Super-Uniform if its cumulative distribution function (CDF) *F* is upper bounded by that of the Uniform distribution (denoted by U[0, 1] below), that is:

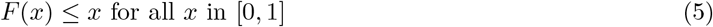

(see e.g. [9, Section 3.3]). Moreover, the closer the *p*-value distribution under the null hypothesis is to U[0, 1], the more powerful the corresponding test is. Condition (5) is satisfied for classical permutation *p*-values such as *p*_*b*_ (with the CDF getting closer to the U[0, 1] distribution as *K* increases), but not for averages of *p*-values like 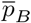. Instead, all the *p*_*b*_ could be correctly aggregated by taking their minimum and correcting the result for multiple testing (Bonferroni aggregation).

If the *p*_*b*_ were independent, then, by the Central Limit Theorem (e.g. [10, Theorem 27.1]), the distribution of 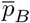 would be asymptotically Gaussian 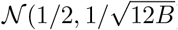 as *B* tends to infinity. This distribution does not verify (5), and therefore tests based on such a distribution are mathematically invalid. In the setting of [1], the *p*_*b*_ are not independent since they have been computed by bootstrapping from one initial sample. However, for small values of *B* (including the choice of *B* = 25 in [1]), the null distribution of (1) deviates only slightly from the asymptotic independence setting. This is illustrated in Figure 1, where the null distribution of (1) is simulated using the parameters chosen in [1]. Details on the simulation and further analyses of the effect of *K* and *B* are included in the SI.

**Figure 1.**
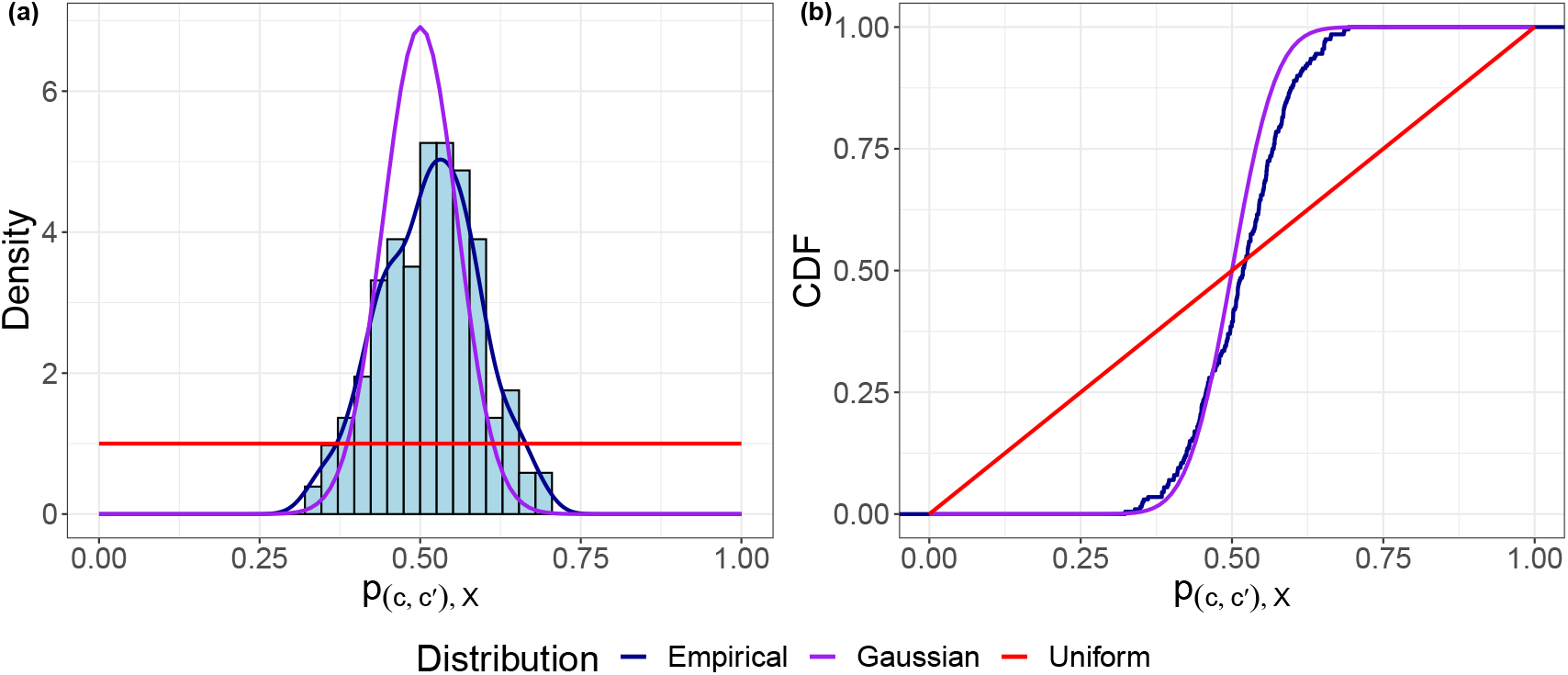
Simulation of the null distribution of *p*_(*c,c′),𝒳*_ for *K* = 200 and *B* = 25, chosen in [1]. Left panel (a): histogram and kernel density estimate. Right panel (b): empirical Cumulative Distribution Function (CDF). Purple lines: asymptotic Gaussian distribution 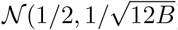; red lines: uniform distribution on [0, 1].

The empirical distribution of *p*_(*c,c′),𝒳*_ presented in Figure 1 does not satisfy Condition (5). Moreover, it is extremely conservative for large values of the statistic realization that is, low *p*-values, yielding an important number of false negatives and thus ignoring substantial differences appearing between the compared samples.

Finally, since the scores *p*_(*c,c′),𝒳*_ are not valid *p*-values, they cannot be incorporated in a multiple testing procedure [11]. In particular, the Benjamini-Hochberg procedure [12] used in [1] needs the *p*-values to be Super-Uniform under the null hypothesis to control the False Discovery Rate (FDR). Consequently, using and adjusting (1) for multiplicity will yield misleading analyses of the overall behaviour of all the null hypotheses and therefore, inaccurate results when the specificities of individual amino acids are studied *a posteriori*.

### Density estimates and reduced sample sizes

Beyond the above-mentioned methodological issues, the approach proposed in [1] presents several practical limitations. It needs, on the one hand, a substantial reduction of sample sizes, which may imply an important loss of information in some cases and thus a substantial power reduction. Indeed, the maximum sample size in [1] was set to *N*_max_ = 200, whereas, for instance, the mean sample size for *α*-helical conformations was 724 and only 18% of the samples had sizes below *N*_max_. On the other hand, it requires a prior parametric estimation of the underlying densities, whose parameters would need to be optimized. In [1], the authors opted to fix the same bandwidth for all comparisons. However, too small bandwidths can lead to density undersmoothing, especially for small sample sizes. This would yield biased kernel estimates whose comparison might lead to false positives.

## Goodness-of-fit between codon-specific (*ϕ, ψ*) distributions

In our recent work [13], we defined two two-sample goodness-of-fit tests for probability distributions supported on the two-dimensional flat torus, in order to study local changes on polypeptide backbone conformations. Both approaches are non-parametric and they use the information provided by entire datasets. The test statistic is based on the 2-Wasserstein distance, which integrates the geometry of the underlying space and provides strong theoretical guarantees and attractive empirical performance [14]. Here, we implemented the first of the testing procedures defined in [13], called *N*_*g*_-geod, to detect differences between the codon-specific Ramachandran plots provided in [1]. For each amino acid, we tested all the pairwise differences between the (*ϕ, ψ*) distributions for all pairs of synonymous codons. As in [1], we kept data points where codons were unambiguously assigned (whose codon scores equal one). Similarly, we repeated the same redundancy filtering based on averaging (*ϕ, ψ*) points with identical Uniprot ID and sequence position. Note that an alternative filtering approach, which kept every first redundant point in the dataset, produced similar results. To facilitate the comparison with the results in [1], we kept only pairs of samples with sizes *n, m* ≥ 30 and we classified all conformations according to their secondary structure according to DSSP [15]: extended strand (E) and *α*-helix (H). We also performed the analysis for all the conformations not belonging to any of these two classes, which we named *Others*. As in [1], the Benjamini-Hochberg multiplicity correction [12] was performed to the computed *p*-values. When representing the Empirical Cumulative Distribution Function (ECDF) of the *p*-values, points laying above the line of slope 1*/α* are considered rejections for a target FDR of *α* according to the BH procedure. The results are presented in Figure 2(a).

**Figure 2.**
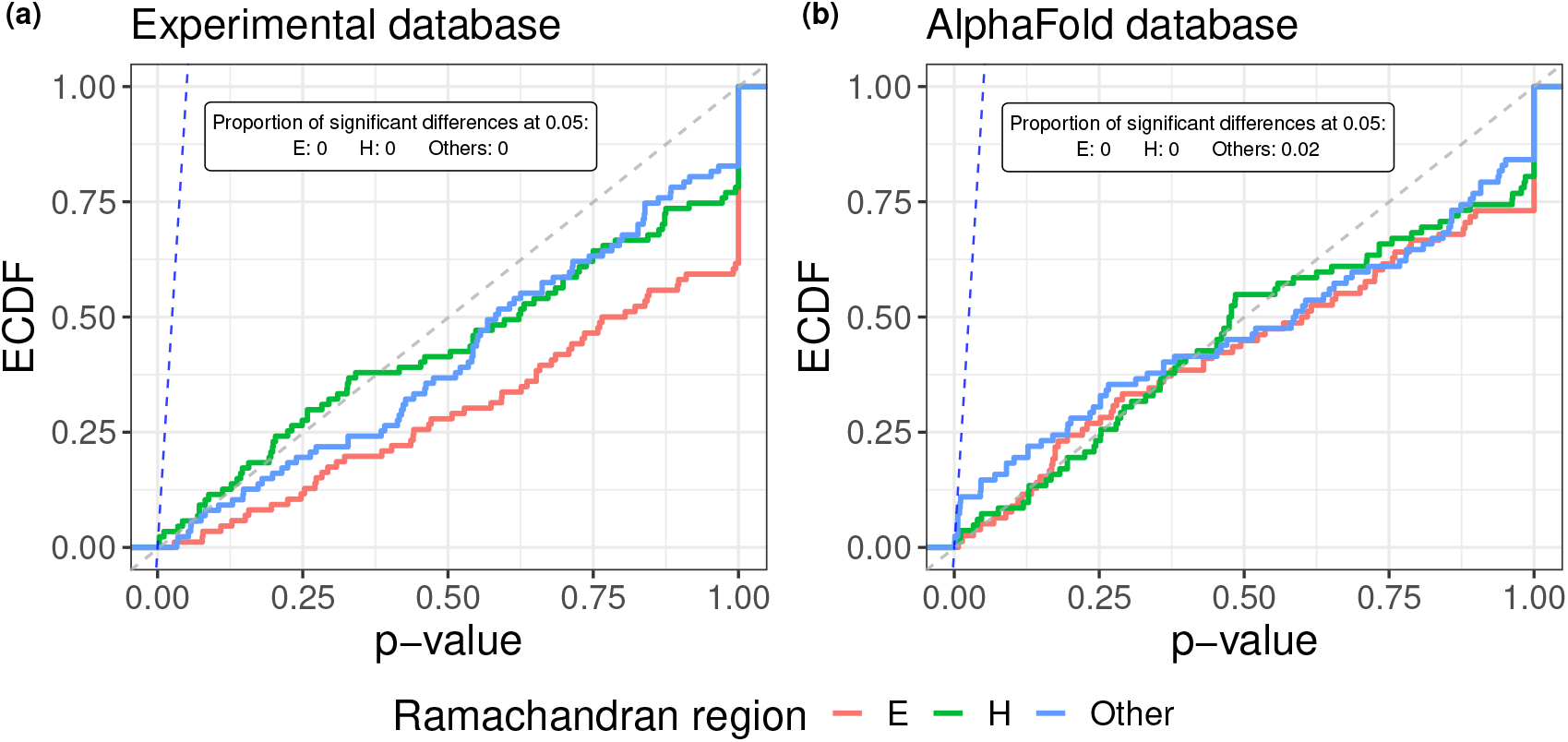
Empirical cumulative distribution function (ECDF) of corrected *p*-values corresponding to testing the equality of (*ϕ, ψ*) distribution pairs corresponding to different synonymous codons in the (a) experimental and (b) AlphaFold database, for conformations in extended strand (E, red), *α*-helix (H, green) and other (*Others*, blue) secondary structures. The dashed blue line of slope 1*/α* corresponds to a target FDR set to *α* = 0.05 for the Benjamini-Hochberg (BH) correction, determining the proportion of rejections among each set of tested hypotheses. The dashed gray line represents the CDF of a Uniform distribution.

The *p*-value distributions presented in Figure 2(a) indicate that, for all the tested hypotheses, differences between codon-specific Ramachandran plots are not significant at level *α* = 0.05. Indeed, the three depicted ECDF lay below the line of slope 1*/α* and therefore no rejections are produced for a target FDR of *α*. Note that our results for *α*-helical structures agree with those presented in [1], for which no significant discrepancies were retrieved. However, results for E structures strongly contradict those in [1], for which significant conformational differences were found for 66% of the synonymous codon pairs tested. Therefore, the main conclusion in [1] is firmly refuted when an appropriate statistical approach is used and, as a consequence, no significant effect of the translated codon identity on the amino-acid backbone conformation can be inferred from the current data.

The discrepancies between the two analyses are most probably due to the above-discussed incorrectness of the methods applied in the original study and, especially, from the potential use of biased density estimates to describe small (*ϕ, ψ*) samples. Indeed, when we looked at the codon pairs whose (*ϕ, ψ*) distributions were found significantly different in [1], we observed that their sample sizes concentrated around the smallest values in the dataset (see Figure 3). This correspondence between significant differences and small sample sizes is counterintuitive for a well-defined statistical test. When the sample size is small and there is limited information about the underlying distribution, the null hypothesis is not rejected unless the evidence against it is very strong. Similarly, *p*-values closer to zero were often found for larger sample sizes, where the statistical power (the test’s ability to detect differences) is higher. This phenomenon, which is depicted in Figure 3 for E structures, suggested that false positives might be appearing due to misleading comparisons of small (*ϕ, ψ*) samples. As we discussed in the previous section, this may be caused by undersmoothed kernel density estimates computed with too small bandwidths for that setting.

**Figure 3.**
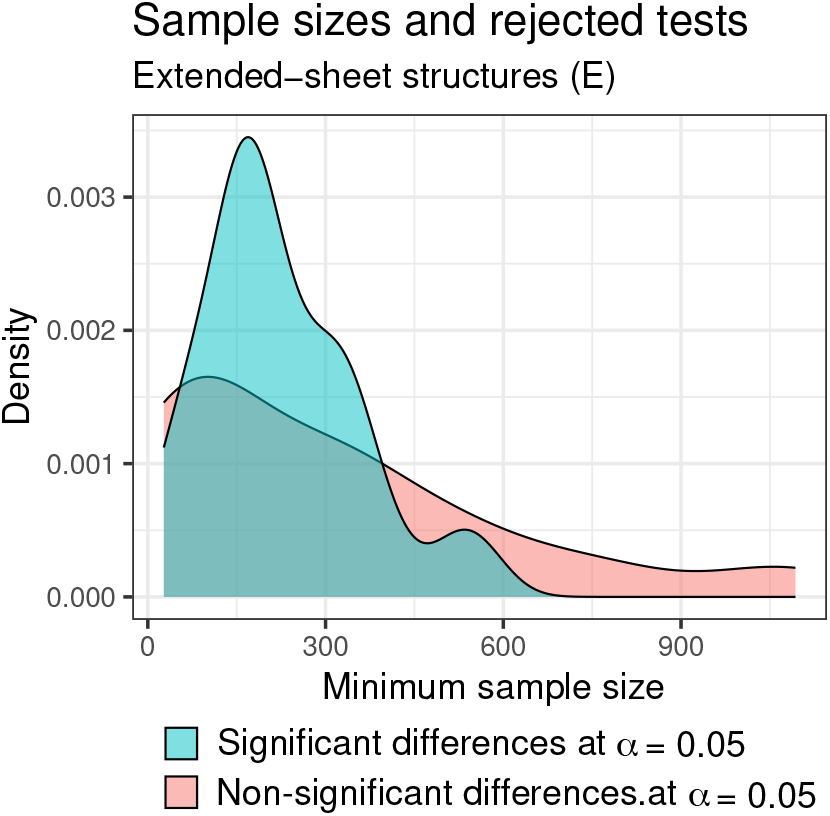
Distribution (kernel density estimates) of the minimum sample size for the codon pairs in the dataset provided in [1], after aggregating redundant data points and removing those with ambiguous codon assignment. Groups correspond to codon pairs for which differences between their codon-specific (*ϕ, ψ*) distributions were found significant (blue) and non-significant (red) at level *α* = 0.05 in [1].

## Additional analyses confirm the robustness of our observations

To validate our conclusions, we performed additional analyses considering different settings. First, we used structures extracted from the AlphaFold Database [7, 8] for the same set of sequences. Note that only residues with pLDDT values larger than 90 were collected for the analysis. Results, depicted in Figure 2(b), qualitatively match those in Figure 2(a) and therefore support the aforementioned conclusions.

The results presented above, as those of Rosenberg *et al*., were based on the structural classification provided by DSSP [15]. We performed the same analyses using a less restrictive classification of structural classes, only considering conformational regions on the Ramachandran space based on non-overlapping angular intervals and disregarding the formation of hydrogen bonds [16, 17]. The corresponding results (see SI) showed that differences between codon-specific Ramachandran plots were statistically non-significant.

Finally, we assessed whether the consideration of the nearest neighbors effects on (*ϕ, ψ*) distributions may alter our results. Indeed, the invalidity of Flory’s Isolated Pair Hypothesis [18] and the interdependence of neighbor effects have been demonstrated in several experimental [19, 20] and theoretical/computational studies [21, 22]. The consideration of neighboring residues is particularly relevant here, as the dataset in [1] exhibits important differences in the proportion of left and right neighboring amino acid types among synonymous codons (see Figure S3). After repeating the same analysis but fixing the identities of left and right neighbors, the overall conclusions remain the same. Results, presented in the SI, show once again that differences between codon-specific Ramachandran plots are non-significant for all secondary structure types.

## Concluding remarks

The work of Rosenberg *et al*. introduced a new paradigm in biology: the nature of the codon influences the (*ϕ, ψ*) angles of protein secondary structures. While the correlation between synonymous codons and secondary structure in the translated proteins is a well known phenomenon [4–6], differences at the (*ϕ, ψ*) level for the most populated conformational states emerged as an intriguing and controversial observation [23]. The conclusions reached from their work could have major impact on one of the paradigms of structural biology, which should shift from protein-sequence to DNA-sequence structure encoding, an information that is not currently stored in structural databases.

With the present study, we have demonstrated the incorrectness of the statistical methodology proposed in [1] to compare probability distributions supported on the two-dimensional flat torus. This, together with the use of density estimates that are not appropriately tuned for small sample sizes, makes the approach in [1] unsuitable to correctly compare codon-specific Ramachandran plots. When applying our previously developed statistical methodology [13] to the same database, no statistically significant differences between the structure encoded by synonymous codons could be detected. Importantly, we demonstrated that this observation is robust with respect to the origin of the 3D structure, the definition of the structural classes and the type of the flanking residues. Therefore, the ensemble of our results unambiguously show that, based on available data, a significant influence of the codon usage in the distribution of backbone dihedral angles in proteins cannot be inferred.

It is worth mentioning, however, that our results have been derived from a limited set of *Escherichia coli* proteins for which the structure had been experimentally determined, and assuming that the gene used for the production of the protein was the same as in the original organism. We believe that a general understanding of the of codon-specific Ramachandran plots can only be achieved by using extensive structural databases, including the corresponding gene sequence, and applying robust statistical methods, such as the one presented here.

## Software availability

The code to reproduce the analyses presented here is available at https://github.com/gonzalez-delgado/synco. The two testing procedures defined in [13] for assessing differences between (*ϕ, ψ*) distributions are implemented in the R package torustest, available at https://github.com/gonzalez-delgado/torustest.

## Author contributions

All the authors designed the studies, interpreted the results and wrote the manuscript; J.G. developed all the computational methods and performed the analyses; J.G. and P.N. carried out the theoretical analyses.

## Competing interests

The authors declare no competing interests.

## Notes

https://github.com/gonzalez-delgado/synco

### Competing Interest Statement

The authors have declared no competing interest.

### Summary of Updates

The dataset used in the previous version was not fully filtered to avoid redundancy. Analyses are repeated using a well-filtered dataset.

