## Supplementary Information (SI) for "The dependence of the amino acid backbone conformation on the translated synonymous codon is not statistically significant"

### A The null distribution of $p$-values is not super-uniform

This section is devoted to prove the technical results stated in the main text about the null distribution of the $p$-values defined in [1], in particular their non-super-uniformity. Keeping the notation of [1], for each pair of bootstrapped samples, we can define the quantity $p_b = (1 + n_b)/(1 + K)$ as a $p$-value for the $b$-th permutation test, with $b = 1, \dots, B$. Each of the $p_b$ follows a super-uniform distribution under the null hypothesis and thus it can be used to define a suitable statistical test accepting correction for multiplicity. Below, we show that the $p$-values defined in [1] are approximately the empirical mean of all the $p_b$, where the approximation error vanishes when the number of permutations $K$ increases.

**Proposition A.1.** *Let $p_{(c,c'),\mathcal{X}}$ be the p-value defined in [1] for a given null hypothesis $H_{0,(c,c')|\mathcal{X}}$, computed as detailed in [1] through $B$ bootstrapped samples with $K$ permutations each. Let $p_b$ denote the p-value for the b-th permutation test for $b = 1, \dots, B$, and $\overline{p}_B$ be the empirical mean of $(p_b)_{1\leq b\leq B}$. Then for any $K > 0$, it holds that:*

$$0 \leq \overline{p}_B - p_{(c,c'),\mathcal{X}} \leq \frac{1}{K} \,. \tag{1}$$

*Proof.* Recalling that the $p$-value for the $b$-th permutation test is $p_b = \frac{n_b+1}{K+1}$, we have

$$p_{(c,c'),\mathcal{X}} = \frac{1 + \sum_{b=1}^{B} n_b}{1 + BK} = \frac{1 + \sum_{b=1}^{B}(p_b(K+1) - 1)}{1 + BK} = \frac{(K+1)\sum_{b=1}^{B} p_b - (B-1)}{1 + BK} \tag{2}$$

$$= \frac{B(K+1)\overline{p}_B - (B-1)}{1 + BK} \,. \tag{3}$$

Therefore, we obtain

$$p_{(c,c'),\mathcal{X}} - \overline{p}_B = \frac{\overline{p}_B(B(K+1) - (1 + BK)) - (B-1)}{1 + BK} = \frac{(B-1)(\overline{p}_B - 1)}{1 + BK} \,. \tag{4}$$

Since $0 \leq p_b \leq 1$ for all $b$, we have $0 \leq \overline{p}_B \leq 1$ as well, so that

$$0 \leq \overline{p}_B - p_{(c,c'),\mathcal{X}} \leq \frac{(B-1)}{1 + BK} \leq \frac{1}{K} \,, \tag{5}$$

where the last inequality holds for any $B$ and $K$ since $(B-1)K \leq 1 + BK$. □

The next step is to show that the empirical mean of two or more uniformly distributed random variables is not super-uniform. As noted in the main text, this empirical mean is asymptotically Gaussian if the variables are independent. The difficulty here is to prove the result without assuming independence, in order for it to be relevant for the study of $p_{(c,c'),\mathcal{X}}$ . We first recall the definition of super-uniformity and give a useful equivalent statement in Remark A.1. Then, Proposition A.2 proves the non-uniformity of $\overline{p}_B$.

**Definition A.1** (Super-uniformity)**.** *A random variable $X$ taking values in $[0,1]$ is said to be super-uniform if it is stochastically greater than a uniform random variable or, in other words, if*

$$\mathbb{P}(X \leq t) \leq t \quad \forall t \in [0,1]\,. \tag{6}$$

**Remark A.1.** *As a consequence of Theorem 1 in [2], a random variable $X$ taking values in $[0,1]$ is super-uniform if and only if*

$$\mathbb{E}(u(U)) \leq \mathbb{E}(u(X)) \quad \textit{for all non-decreasing function } u\,, \tag{7}$$

*where $U$ denotes a random variable uniformly distributed in $[0,1]$.*

**Proposition A.2.** *Let $U_1, \dots, U_n$ be $n$ real-valued random variables uniformly distributed on $[0,1]$. For all $n \geq 2$, their empirical mean $\overline{U_n} = \frac{1}{n}\sum_{i=1}^{n} U_i$ is not super-uniform.*

*Proof.* Let $U$ be a random variable uniformly distributed in $[0,1]$. As a consequence of Remark A.1, it suffices to find a non-decreasing function $u$ such that $\mathbb{E}(u(U)) > \mathbb{E}(u(\overline{U_n}))$ for all $n \geq 2$. Let $u : [0,1] \longrightarrow [0,1]$ be such that $u(t) = t^2$ for all $t \in [0,1]$. Then, as $\mathbb{E}(\overline{U_n}) = \mathbb{E}(U)$ and $\mathbb{E}(X^2) = \mathrm{Var}(X) + \mathbb{E}(X)^2$ for any real-valued random variable $X$, it suffices to prove that

$$\mathrm{Var}\left(\overline{U_n}\right) < \mathrm{Var}(U) = \frac{1}{12} \quad \forall\, n \geq 2\,. \tag{8}$$

First, we have

$$\mathrm{Var}\left(\overline{U_n}\right) = \frac{1}{n^2}\mathrm{Var}\left(\sum_{i=1}^{n} U_i\right) = \frac{1}{n^2}\left[\sum_{i=1}^{n}\mathrm{Var}(U_i) + 2\sum_{i<j}\mathrm{Cov}(U_i,U_j)\right] = \tag{9}$$

$$\frac{1}{12n} + \frac{2}{n^2}\sum_{i<j}\mathrm{Cov}(U_i,U_j) = \frac{1}{12n} + \frac{1}{n^2}\sum_{i<j}\left(\mathbb{E}(U_iU_j) - \mathbb{E}(U_i)\mathbb{E}(U_j)\right) = \tag{10}$$

$$\frac{1}{12n} + \frac{2}{n^2}\sum_{i<j}\left(\mathbb{E}(U_iU_j) - \frac{1}{4}\right) = \frac{1}{12n} - \frac{1}{2n^2}\binom{n}{2} + \frac{2}{n^2}\sum_{i<j}\mathbb{E}(U_iU_j)\,. \tag{11}$$

As the expectation of the product of two random variables defines an inner product on the set of random variables equally supported, we can apply Cauchy–Schwarz inequality and upper bound the last expectation in (11) as

$$\mathbb{E}(U_iU_j) \leq \sqrt{\mathbb{E}(U_i^2)\mathbb{E}(U_j^2)} = \frac{1}{3}\,. \tag{12}$$

However, the maximum $\frac{1}{3}$ is achieved if and only if both random variables are equal. Indeed, an equality in (12) holds if and only if the two variables are linearly dependent [3]. If, what is more, they are identically distributed, linear dependence is equivalent to equality. Consequently, at least one of the pairs $i < j$ must satisfy $\mathbb{E}(U_iU_j) < \frac{1}{3}$ or, on the contrary, we would have $U_1 = \cdots = U_n$, contradicting the hypothesis $n \geq 2$. Therefore, we can upper bound (11) as

$$\mathrm{Var}(\overline{U_n}) < \frac{1}{12n} - \frac{1}{2n^2} + \frac{2}{3n^2}\binom{n}{2} = \frac{1}{12n} + \frac{1}{6n^2}\binom{n}{2} = \frac{1}{12} \quad \forall\, n \geq 2\,, \tag{13}$$

which concludes the proof. □

### B Numerical study of $p$-value null distribution

In this section, we illustrate the behaviour of the non-uniform $p$-values $p_{(c,c'),\mathcal{X}}$ under the null hypothesis. As explained in the main text, the $B$ individual $p$-values $p_b$, $b = 1, \ldots, B$, are not independent as they are computed by bootstrapping from one initial sample. If, on the contrary, the $p_b$ were computed from independent samples, the empirical mean $\overline{p}_B$ would converge in distribution to a Gaussian (Theorem 27.1 in [4]):

$$\sqrt{12B}\left(\overline{p}_B - \frac{1}{2}\right) \xrightarrow[B\to\infty]{\mathcal{D}} \mathcal{N}(0,1)\,. \tag{indep}$$

We aim at analysing how the dependence induced by bootstrapping alters the asymptotic distribution of (indep), as well as the effect of the number $B$ of bootstrap iterations and the number $K$ of permutations for each individual test. We simulated the distribution of $p_{(c,c'),\mathcal{X}}$ under the null hypothesis following the algorithm detailed in [1] (see *Full procedure* in Methods Section, end of p. 7). The original samples were drawn from a uniform distribution and had fixed sizes $N = 2000$. Bootstrapped samples were extracted with size $N_{\max} = 200$. For the independence scenario, we replaced the bootstrapped samples by new equally sized samples drawn from a uniform distribution. As the explicit form of the test statistic is not provided in [1], we used the Wilcoxon statistic to illustrate the behaviour of $p_{(c,c'),\mathcal{X}}$. For each pair of values of $K, B$, the null distribution of $p_{(c,c'),\mathcal{X}}$ was simulated with 200 Monte Carlo iterations. Results are presented in Figure S1, where the empirical distribution is compared to the asymptotic independence scenario (indep).

The first row in Figure S1 shows the null distribution of $p_{(c,c'),\mathcal{X}}$ if samples are not bootstrapped but drawn independently at each iteration $b = 1, \ldots, B$. The encountered empirical distribution matches the Gaussian (indep) more faithfully as $K$ increases, which was expected as the difference $\left\|p_{(c,c'),\mathcal{X}} - \overline{p}_B\right\|$ is upper bounded by $1/K$. In the same way, we should expect that the simulated $p_{(c,c'),\mathcal{X}}$ distributions are closer to the real (and unknown for this dependency scenario) null distribution of $\overline{p}_B$ when moving from the left to the right column in Figure S1. When samples are bootstrapped as in [1], dependency between the $p_b$ appears and as $B$ increases (from the second to the last row in Figure S1) values deviate from the independence scenario (indep). When $B$ remains small (as for $B = 25$, the value chosen in [1]), the deviation from (indep) is slight. This can be explained as $N_{\max} \ll N$, and bootstrapping few times samples with small size compared to the one of the original sample is close to drawn samples independently from the entire population. As $B$ increases, so does the dependency between the individual $p$-values. This dilates the empirical distribution of $p_{(c,c'),\mathcal{X}}$ and extends the difference to (indep). A similar phenomenon was observed in [5] when studying the effect of unobserved covariates on the null distribution of $p$-values.

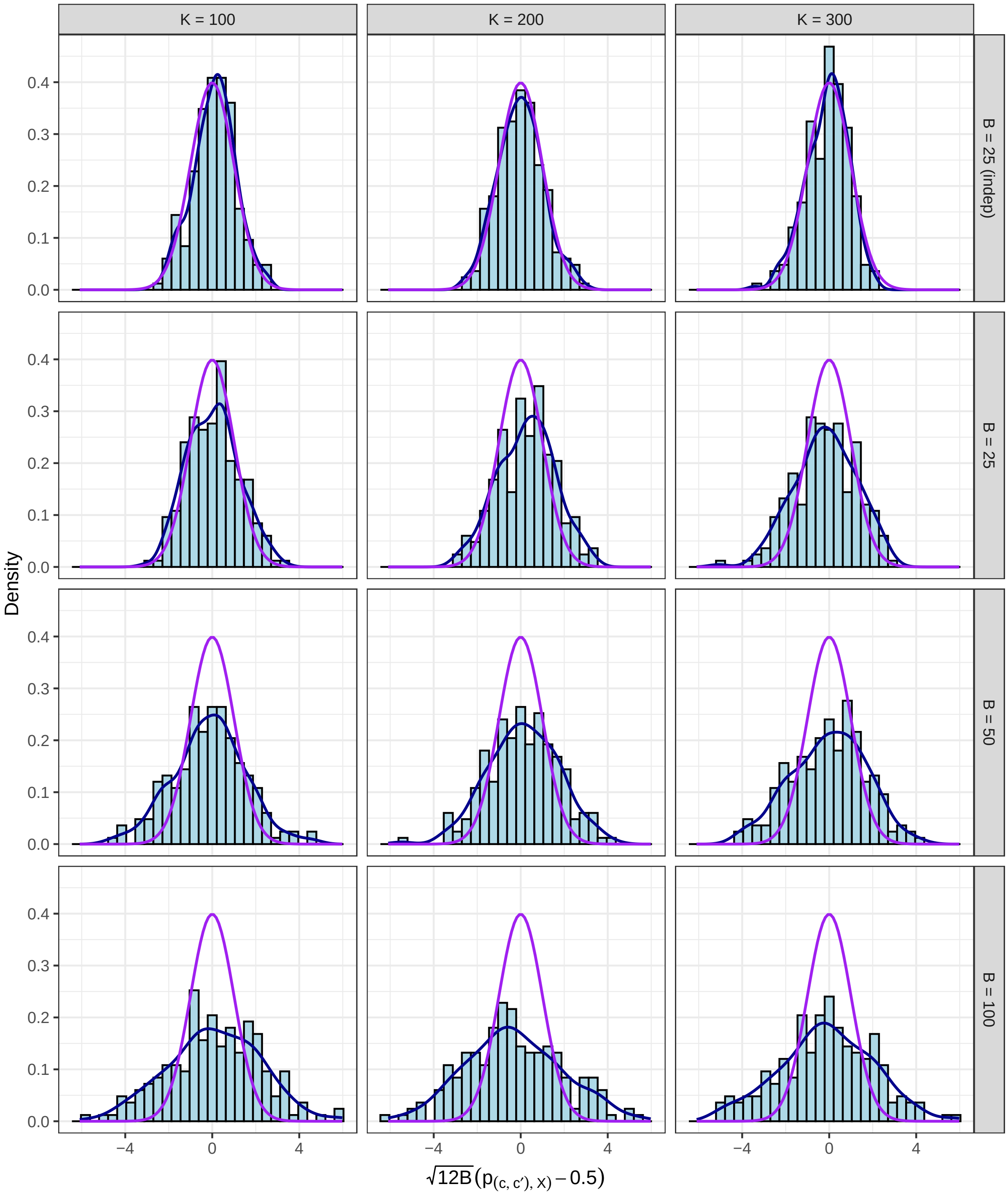


Figure S1: Simulation of the null distribution of $p_{(c,c'),\mathcal{X}}$ for different values of parameters $K$ (in columns) and $B$ (in rows). The first row corresponds to the independence scenario (indep), whose asymptotic standard Gaussian density is depicted in purple in all cases. The blue line corresponds to the non-parametric kernel density estimate of the encountered empirical distribution. Note that, for the sake of comparison to (indep), the presented $p$-values have been re-scaled as $\sqrt{12B}(p_{(c,c'),\mathcal{X}} - 0.5)$.

### C Structural classification as non-overlapping regions of the Ramachandran space

We repeated the analysis presented in Figure 2 of the manuscript using a different structural classification, simply based on non-overlapping regions on the Ramachandran space:

$$\mathcal{A} = (-180°, 0°] \times (-120°, 50°], \quad \mathcal{B} = (-180°, 0°] \times (-50°, 240°]. \tag{14}$$

Note that classes $\mathcal{A}$ and $\mathcal{B}$ are not limited to $\alpha$-helices and extended strands. For instance, poly-l-proline type II (PPII) structures are included in $\mathcal{B}$. Moreover, a substantial number of conformations that were not classified as $\alpha$-helical (H) or extended stand (E) by DSSP (named 'Others' in Figure 1) belong now to the $\mathcal{A}$ or $\mathcal{B}$ classes. More precisely, 23.73% and 17,82% of 'Others' conformations are now contained in $\mathcal{A}$ or $\mathcal{B}$, respectively. Results of this analysis are presented in Figure S2, and discussed in the main text.

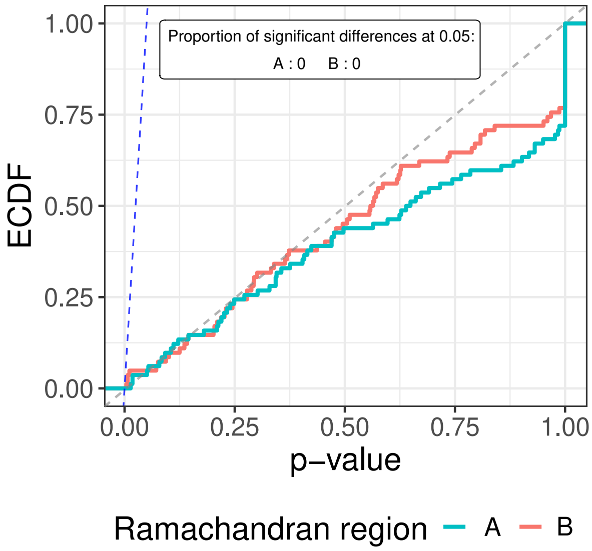


Figure S2: Empirical cumulative distribution function (ECDF) of corrected $p$-values corresponding to testing the equality of $(\phi, \psi)$ distribution pairs corresponding to different synonymous codons, for conformations in $\mathcal{A}$ and $\mathcal{B}$ classes, defined by the angular intervals (14). The dashed blue line of slope $1/\alpha$ indicates a target FDR set to $\alpha = 0.05$ for the Benjamini-Hochberg (BH) correction, determining the proportion of rejections among each set of tested hypotheses. The dashed gray line represents the CDF of a Uniform distribution.

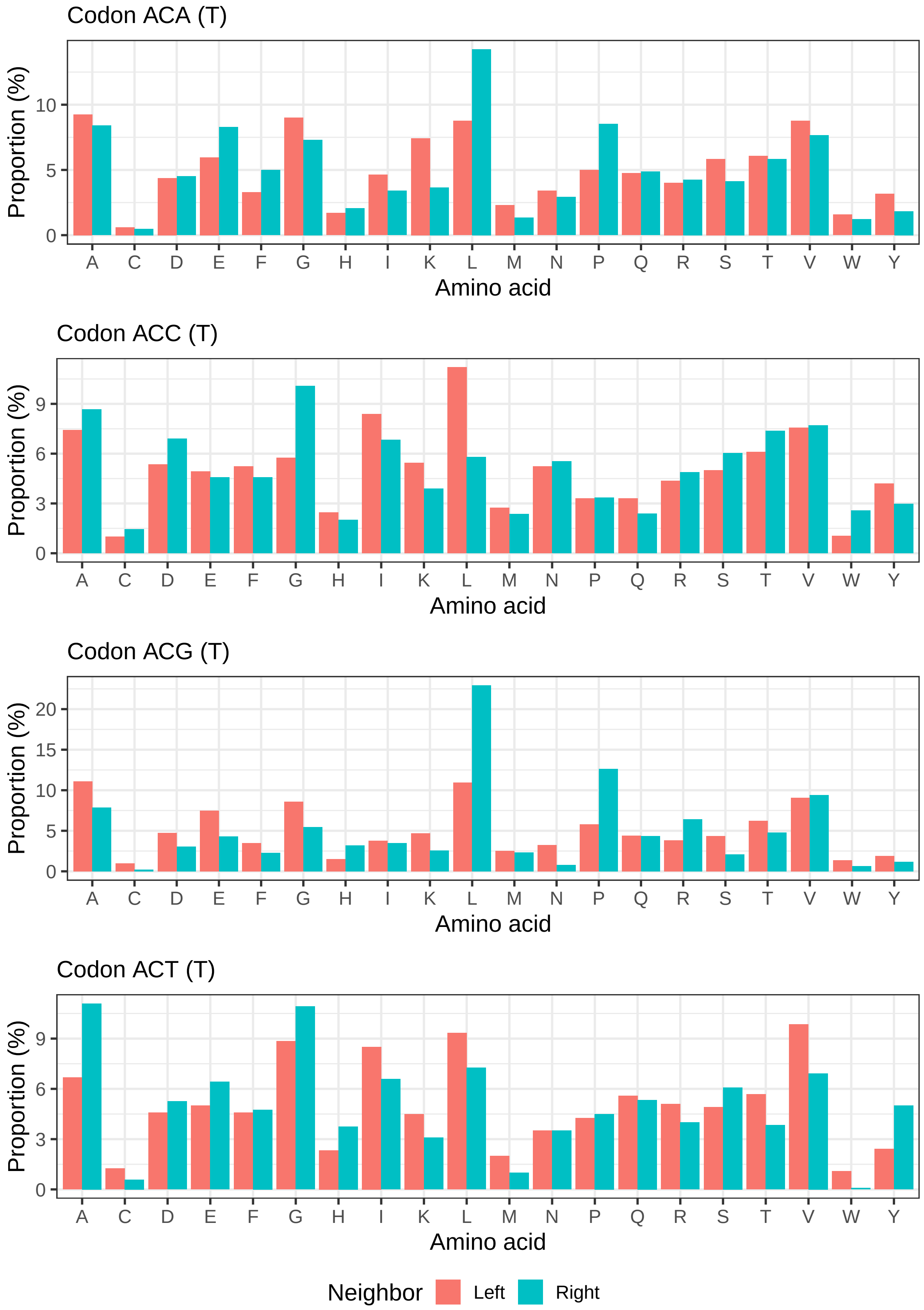


Figure S3: Proportions (in percentage) of left and right neighboring amino acid types for the four synonymous codons of threonine in the dataset provided in [1], after aggregating redundant data points and removing those with ambiguous codon assignments.

### D  Tripeptide-specific $(\phi, \psi)$ distribution analysis

The relevance of the nearest neighbor effects on the conformational preferences of an amino acid residue, in terms of $(\phi, \psi)$ distribution, has been demonstrated using experimental and computational approaches. Nevertheless, these effects have not been taken into account when defining the codon-specific distributions in [1]. As discussed in the main text, the proportion of left and right neighboring amino acid types substantially differs among synonymous codons. This is illustrated in Figure S3 for threonine. The behavior is similar for all amino acid types. Consequently, neighbor effects do not equally influence each $(\phi, \psi)$ distribution corresponding to different synonymous codons, introducing a bias that remains uncontrolled in [1]. However, when repeating the analyses by considering codon-specific Ramachandran plots for triplets of amino acids, the overall conclusions do not change. The corresponding results are presented in Figure S4. As sample sizes considerably decreased, we kept codon pairs having more than 10 points in both samples.

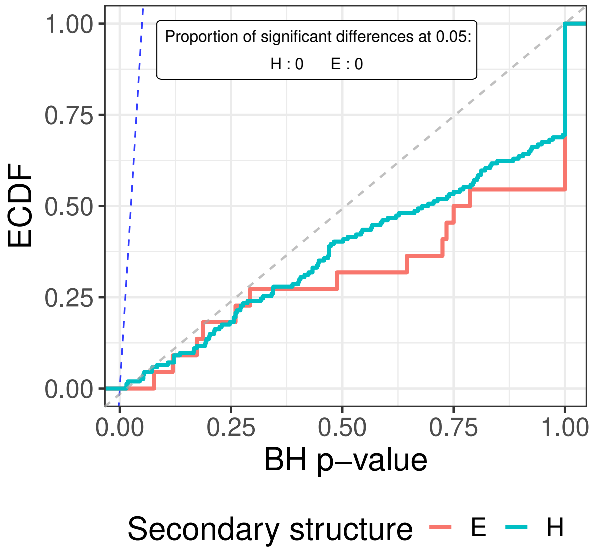


Figure S4: Empirical cumulative distribution function (ECDF) of corrected $p$-values corresponding to testing the equality of $(\phi, \psi)$ distribution pairs corresponding to different synonymous codons and identical left and right neighbors, for conformations in extended strand (E) and $\alpha$-helix secondary structures. The dashed blue line of slope $1/\alpha$ indicates a target FDR set to $\alpha = 0.05$ for the Benjamini-Hochberg (BH) correction, determining the proportion of rejections among each set of tested hypotheses. The dashed gray line represents the CDF of a Uniform distribution.
